# SimulateCNVs: a novel software application for simulating CNVs in WES and WGS data

**DOI:** 10.1101/407486

**Authors:** Yue Xing, Alan R. Dabney, Xiao Li, Claudio Casola

## Abstract

**Summary:** SimulateCNVs is a fast and novel software application for simulating CNVs in WES and WGS data. Current NGS simulators cannot simulate rearranged genomes and their alignment files for WES data and are not easy to use. SimulateCNVs is the first software application that can simulate CNVs in both WES and WGS data, is easy to install, has many unique features, and can output rearranged genomes, short reads and bam files in a single command.

**Availability:** SimulateCNVs is freely available from https://yjulyxing.github.io/

---

Copy Number Variants (CNVs) are insertions and deletions of 1 kb or larger in a genome, a type of structural variant that has been associated with phenotypic changes, including disease in human (Feuk, et al., 2006). Increasingly, next-generation sequencing (NGS) data are used to detect CNVs. Many software applications have been developed to detect CNVs using either whole-genome sequencing (WGS) [2] or whole-exome sequencing (WES) [3] data. Most of these methods are based on read depth, although approaches relying on read pairs distance, detection of split reads or *de novo* genome assemblies have also been developed for WGS data [2]. Combinations of some of these approaches have also been incorporated in CNV detection programs [2]. Overall, there is poor agreement between methods due to a variety of issues. Simulations of copy-number variants in WGS and WES datasets allow comprehensive comparisons of the operating characteristics of existing and novel CNV detection methods.

There are several existing software applications for simulating CNVs and other structural variants in NGS data, including RSVSim (Bartenhagen and Dugas, 2013), SCNVSim (Qin, et al., 2015), Pysim-sv (Xia, et al., 2017), SVsim (https://github.com/GregoryFaust/SVsim) and SInC (Pattnaik, et al., 2014). However, the current versions of these applications are not easy to use and have many limitations when used for CNV simulation with WES data. In WES data, specific regions of a reference genome, called “target regions”, are captured and sequenced (Goh and Choi, 2012). To reproduce a realistic distribution of structural variants, a CNV simulator for WES data should generate variants that overlap partly or entirely with one or multiple target regions (see Supplemental File 1, Supplementary Figure 1). Genomic coordinates of these regions should be included in a separate file. Short reads are then produced from both control (same as reference genome) and test (with simulated CNVs) genomes and aligned back to the reference genome. Insertions and deletions in the test genome appear as increased read coverage or lack of coverage, respectively, in the reads alignment file (Supplemental File 1, Supplementary Figure 1). Some of the above methods can simulate CNVs in selected regions, but they lack the capacity to rearrange the target regions in the rearranged genome according to simulated CNVs (Supplemental File 1, Supplementary Figure 1). Therefore, these methods cannot be used to simulate WES datasets, because they do not provide the locations of the rearranged target regions necessary to synthesize short reads. Previous methods also lack the capacity to produce CNVs that overlap partly or entirely with one or multiple target regions. VarSimLab is the only current program specifically designed to simulate CNVs with WES data (https://varsimlab.readthedocs.io/en/latest/index.html, (Zare, et al., 2017); previously called CNV-Sim: https://nabavilab.github.io/CNV-Sim/). VarSimLab shows several limitations that make its simulated datasets unrealistic and of limited value. Briefly, VarSimLab cannot accommodate varying degrees of CNV overlap with target regions, cannot simulate pooled samples with a common control, only accepts one chromosome at a time, and simulates CNVs that overlap greatly with one another; for more on VarSimLab limitations, see Supplemental File 2.

Here, we introduce SimulateCNVs, a novel tool that overcomes known limitations of existing CNV simulation tools, as a software application for simulating CNV datasets to facilitate comparison of the performance of different CNV detection methods. SimulateCNVs is python-based, can simulate CNVs in either the WGS or WES context, and can be easily installed and used on Linux systems. SimulateCNVs is particularly well suited to simulating CNVs in WES data. Users can use a single command/step to generate rearranged genomes, short reads and bam files for multiple test samples, with the control being the original genome. SimulateCNVs outputs rearranged genomes, with non-overlapping CNVs that are either designed by the user or randomly generated, and target regions for sequencing. Additionally, it accepts genomes with multiple chromosomes. Users have the option to then employ SimulateCNVs to generate short read files by simulating Illumina single or paired-end sequencing for test and control samples. This step includes custom codes written to utilize the WGS short read simulator ART_illumina (Huang, et al., 2011) to sequence the target regions/genome. Finally, users can choose to simulate bam files from short read files for tests and control. This step is a routine pipeline utilizing BWA (Li, 2013), samtools (Li, 2011; Li, et al., 2009), picard (v2.15.0, https://broadinstitute.github.io/picard/) and GATK (McKenna, et al., 2010).

SimulateCNVs also includes the following optional features:

1. CNVs will not overlap with any gap regions (stretches of “Ns”). An optional script is also provided for replacing gap regions with nucleotides randomly.
2. Set the minimum distance in bp between CNVs.
3. Custom management of overlap between simulated CNVs and user-specified target regions. Can choose the minimum number of overlapping base pairs.
4. Set a distance so that target regions closer than that distance can be connected in a single region for sequencing. Users can also choose to sequence up-and downstream of the target regions.
5. CNV start point and length can be chosen randomly, from a distribution or read in from a file providing start, end and copy number of each CNV. Length can also be read from a file solely containing lengths and number of CNVs of each length.
6. If choosing to generate CNVs randomly, users can choose to generate a specified number of CNVs on each chromosome/scaffold, or specify a total CNV number for the whole genome. In the latter case, the number of CNVs will be proportional to the length of chromosomes/scaffolds, with at least 1 CNV generated in the shortest chromosome/scaffold.
7. If a chromosome is too short or there are too many gap regions to accommodate the number of CNVs requested, the program will report the actual number of CNVs generated instead of the planned amount and continue to the next chromosome. This feature prevents program-killing errors like those thrown in such cases by other applications.
8. User-specified proportion of CNVs to be insertions/deletions.

To run the program, the user is required to provide a genome file in fasta format. For WES simulation, a file of target regions is also required. Users specify if gap regions should be avoided for CNV simulation and the minimum length between any two CNVs. Users then specify the method for CNV simulation; for example, number, distribution, range of length and copy numbers of CNVs, and proportion of insertions. For WES simulation, users choose the minimum base pairs of target regions to be overlapping with CNVs and specify how many CNVs overlapping with and outside of target regions should be simulated. Users specify output directory, output name, number of test samples to be simulated and if control should be given. If short reads are required to be simulated, users choose the maximum distance for target regions to be connected for sequencing, in addition to distance up- and down-stream of target regions to be sequenced. Users can specify additional sequencing parameters including fold coverage, single or paired-end sequencing, base quality, etc. Finally, if users choose to simulate bam files with indexes, the local absolute path to picard and GATK must be provided; samtools and BWA are also required to be installed.

The computation time for various genomes is shown in Supplemental File 1, Supplementary Table 1.

## Acknowledgements

The high performance research computing system Ada at Texas A&M University was used for testing the software application.

